# Mechanism of the CBM35 domain in assisting catalysis by Ape1, a *Campylobacter jejuni* O-acetyl esterase

**DOI:** 10.1101/2021.09.23.461501

**Authors:** Chang Sheng-Huei Lin, Ian Y. Yen, Anson C. K. Chan, Michael E. P. Murphy

## Abstract

Peptidoglycan (PG) is *O*-acetylated by bacteria to resist killing by host lysozyme. During PG turnover, however, deacetylation is a prerequisite for glycan strand hydrolysis by lytic transglycosylases. Ape1, a de-*O*-acetylase from *Campylobacter jejuni*, is a bi-modular protein composed of an SGNH hydrolase domain and a CBM35 domain. The conserved Asp-His-Ser catalytic triad in the SGNH hydrolase domain confers enzymatic activity. The PG binding mode and function of the CBM35 domain in de-*O*-acetylation remained unclear. In this paper, we present a 1.8 Å resolution crystal structure of a complex between acetate and Ape1. An active site cleft is formed at the interface of the two domains and two large loops from the CBM35 domain form part of the active site. Site-directed mutagenesis of residues in these loops coupled with activity assays using *p*-nitrophenol acetate indicate the CBM35 loops are required for full catalytic efficiency. Molecular docking of a model *O*-acetylated hexasaccharide PG substrate to Ape1 using HADDOCK suggests the interaction is formed by the active cleft and the saccharide motif of PG. Together, we propose that the active cleft of Ape1 diverges from other SGNH hydrolase members by using the CBM35 loops to assist catalysis. The concave Ape1 active cleft may accommodate the long glycan strands for selecting PG substrates to regulate subsequent biological events.

## Introduction

Bacterial peptidoglycan (PG) is responsible for osmotic stress resistance and cell shape maintenance. The PG polymer consists of linear glycan strands of repeating β-(1-4)-*N*-acetylglucosamine (GlcNAc) and *N*-acetylmuramic acid (MurNAc) units crosslinked by short peptide stems attached to the MurNAc residues. The MurNAc residue of nascent PG in most Gram-negative bacteria is covalently linked to the pentapeptide composed of L-alanine (L-Ala), D-glutamate (D-Glu), *meso*-diaminopimelic acid (*m*-DAP), and two terminal D-alanine (D-Ala) residues. Nascent PG is polymerized to existing PG by transglycosylases, and transpeptidases catalyze 4-3 cross-linkages between (D) *m*-DAP and D-Ala or 3-3 cross-linkages between (L) *m*-DAP and D-Ala of neighboring peptide stems (1). The structure of PG is constantly modified during cell growth and division, as well as to protect against host defenses and antibiotics, and for the assembly of trans-envelope machineries like pili and flagella (2). PG modifications include cleavage of glycosidic bonds by muramidases to reduce glycan chain length, peptide stem cleavage by carboxypeptidases, and peptide cross-linkage cleavage by endopeptidases.

*O*-acetylation at the C6 hydroxyl group of MurNAc adds another level of chemical diversity to mature PG, and serves in a protective role to resist cell lysis by lysozymes produced by mammalian innate immune systems. Pathogens such as *Staphylococcus aureus, Enterococcus faecalis, Listeria monocytogenes* and *Neisseria gonorrhoeae* that contain *O*-acetylated PG are found to be more resistant to lysozymes (3-6). The level of PG *O*-acetylation is determined by enzymes encoded by the O-acetylation of peptidoglycan (OAP) regulon (7). In Gram-positive bacteria, the bi-functional protein OatA translocates cytoplasmic acetyl moieties to the periplasm and acetylates the C6 hydroxyl group of PG MurNAc (8-10). In Gram-negative bacteria, a two-component system, PatA and PatB, performs the activities of OatA (11-13). The OAP regulon encodes a third gene named *ape1*. Ape1 is a periplasmic *O*-acetyl esterase that hydrolyzes *O*-acetylated MurNAc within the PG polymer to produce de-*O*-acetylated MurNAc (14). De-*O*-acetylation by Ape1 is proposed to regulate PG turnover by lytic transglycosylases (LTs) (15), which can only cleave the β-(1-4)-glycosidic bond between de-*O*-acetylated MurNAc and GlcNAc to generate glycan stands with an 1,6-anhydroMurNAc end (16).

*Campylobacter jejuni* is a Gram-negative, helical-shaped, human enteric pathogen. It is a leading cause of bacterial food-borne diarrhea. Infections by *C. jejuni* can lead to autoimmune responses in intestines (e.g., inflammatory bowel disease), joints (e.g., reactive arthritis) and nerves (e.g., Guillain-Barré Syndrome) (17,18). The OAP regulon, particularly the *ape1* gene, contributes to *C. jejuni* pathogenesis (19). An *ape1* deletion strain showed increased PG *O*-acetylation, irregular comma-shaped cell morphologies, and a 30% reduction of wild-type motility on agar plates. The Δ*ape1* mutant strain was significantly impaired in invasion of intestinal INT407 cells and in a chick model post infection showed a 4.4-log CFU/g decrease in the cecum compared to wild-type. Furthermore, Δ*ape1* PG composition showed reduced 1,6-anhydroMurNAc content and elongated glycan strands, supporting the hypothesis that Ape1 activity is an important prerequisite for LT cleavage (19,20). Conversely, deletion of the entire OAP regulon or either *patA* or *patB* in a *C. jejuni* strain showed reduced *O*-acetylation levels in PG and have the same glycan strand length of wild-type (19). These mutant strains did not have defects in motility or chick colonization ability.

Ape1 belongs to the SGNH hydrolase superfamily (7). The catalytic triad consists of Ser, His, and Asp and opposite this triad are Gly and Asn residues that form the oxyanion hole for transition state stabilization in the active site (21,22). Mutagenesis of the *N. gonorrhoeae* Ape1 (NgApe1) homolog demonstrated that these conserved active site residues are required for enzyme activity (14,21,23). NgApe1 has higher specific activity towards *O*-acetylated PG compared to *O*-acetylated xylan (14), suggesting that the enzyme may recognize additional substrate components beyond the acetyl group. The *N. meningitidis* Ape1 (NmApe1) crystal structure contains a catalytic domain with an α/β/α fold of the SGNH superfamily (24). The authors observed that the presence of an acetyl moiety rotates the Ser nucleophile by 90° to be positioned for optimal catalysis and proposed a substrate-induced mechanism as to prevent accidental de-*O*-acetylation. The structure also has a carbohydrate binding module of family 35 (CBM35) in addition to the SGNH domain. The function of such domains in Ape1 is not known. The CBM35 domain is typically found in plant cell wall degrading enzymes and is responsible for guiding glycosidase domains to uronic acid containing substrates (25,26).

In the present study, we determined a 1.8 Å acetate-bound Ape1 crystal structure from *C. jejuni*. We investigated the role of two loops from the CBM35 domain in Ape1 catalysis. Lastly, we performed molecular docking to propose an Ape1-PG binding mode. Together, our results highlight the participation of the previously uncharacterized CBM35 domain in Ape1 catalysis.

## Results

### Structure determination of C. jejuni Ape1

*C. jejuni* Ape1 is a 392 amino acid protein containing a predicted signal peptide at residues 1-21 of the native sequence (*CJJ81176_0638*) **(Figure 1)**. The full-length mature protein (Ape1^22-392^) was crystallized, and despite diffracting to high resolution (better than 1.6 Å), structure determination was impeded by crystal twinning. A truncated construct with residues 41-392 (Ape1^41-392^) was crystallized and its structure was solved to 1.8 Å resolution by single anomalous dispersion (SAD) with selenomethionine-labeled protein (SeMetApe1^41-392^). An initial model was automatically built with Phenix AutoBuild (27), resulting in three partial Ape1 monomers in the asymmetric unit. A continuous model was built for two monomers, with the third monomer incomplete due to disorder.

**Figure 1.**
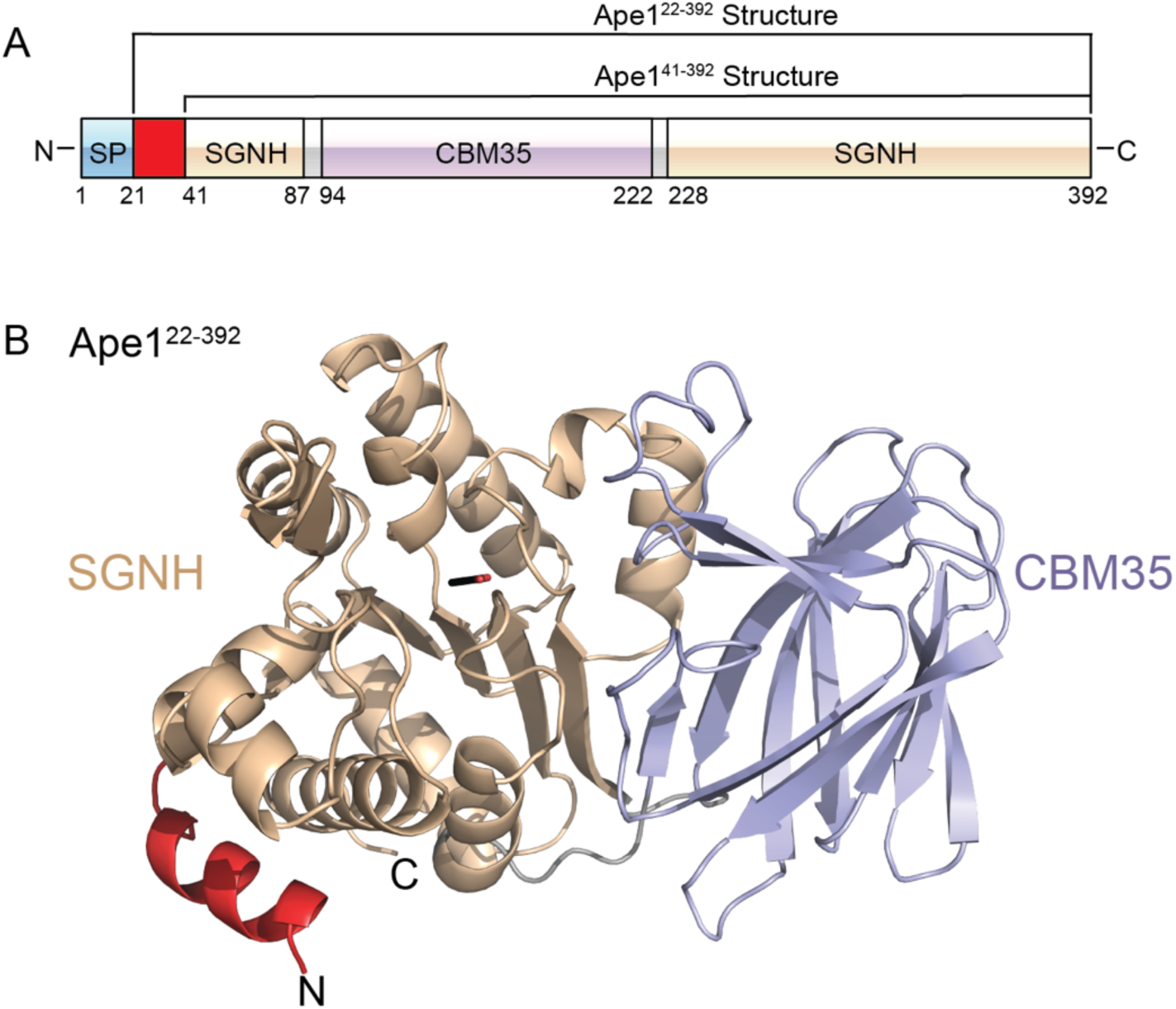
The crystal structure of *C. jejuni* Ape1. (A) Schematic representation of full-length Ape1 (SP = signal peptide). The regions corresponding to recombinant Ape1^41-392^ and Ape1^22-392^ proteins are labeled. (B) The overall structure of acetate-bound Ape1^22-392^ with the N-terminal helix, SGNH and CBM35 domains colored in red, brown and light purple, respectively. The acetate is shown in stick form.

The native Ape1^41-392^ protein structure was solved to 1.8 Å resolution by molecular replacement with one of the complete monomers from the SeMetApe1^41-392^ structure. Three monomers were built and refined to R_work_ and R_free_ values of 0.17 and 0.19, respectively. The final model includes three complete Ape1 monomers (residues 41-392), with each monomer bound to one acetate molecule in the active site. The electron density for the acetate was clearly defined in electron density maps and was refined to an average *B*-factor of 20.7 Å^2^. To explore the oligomeric state, Ape1^41-392^ was analysed by SEC-MALS. In solution, the molecular weight was determined to be 37.3 ± 2.5% kDa, consistent with the predicted molecular weight of the recombinant Ape1 monomer (41.1 kDa).

The Ape1^22-392^ structure was solved by molecular replacement with Ape1^41-392^ as the search model and refined assuming merohedral twinned to R_work_ and R_free_ values of 0.14 and 0.18. The Ape1^22-392^ and Ape1^41-392^ structures are similar, with a RMSD of 0.16 Å over 352 aligned Cα atoms. The Ape1^22-392^ structure also contains three monomers in the asymmetric unit and reveals an additional N-terminal helix **(Figure 1B)**. The N-terminal helix of Ape1^22-392^ shows limited contacts to the rest of the protein and its sequence is not conserved amongst homologs from different species. As the Ape1^41-392^ structure is of higher quality, it was used for all subsequent analyses and will be denoted as CjApe1. All data collection, phasing and refinement statistics are summarized in **Table 1**.

**TABLE 1.** Data collection and model refinement statistics of *C. jejuni* Ape1

|  | Ape1<sup>41-392</sup> | SeMetApe1<sup>41-392</sup> | Ape1<sup>22-392</sup> |
| --- | --- | --- | --- |
| **Data collection<sup>a</sup>** |  |  |  |
| Space group | P3<sub>2</sub> | P3<sub>2</sub> | P3<sub>2</sub> |
| Cell dimensions |  |  |  |
| *a*, *b*, *c* (Å) | 94.9, 94.9, 102.7 | 95.3, 95.3, 103.1 | 95.3, 95.3, 103.0 |
| $\alpha$, $\beta$, $\gamma$ (°) | 90, 90, 120 | 90, 90, 120 | 90, 90, 120 |
| Total reflections | 546678 | 696966 | 271029 |
| No. of unique reflections | 95429 | 111763 | 46459 |
| Wavelength (Å) | 0.97946 | 0.97915 | 0.98011 |
| Resolution (Å) | 50.0–1.80 (1.83–1.80) | 50.0–1.70 (1.73–1.70) | 50.0–2.30 (2.34–2.30) |
| R<sub>merge</sub> | 0.100 (0.798) | 0.087 (0.614) | 0.148 (0.782) |
| *I*/σ*I* | 18.3 (1.7) | 23.0 (2.6) | 13.2 (2.3) |
| CC (1/2) | (0.810) | (0.859) | (0.653) |
| Completeness (%) | 99.7 (99.7) | 97.1 (98.6) | 100 (100) |
| Redundancy | 5.7 (5.1) | 6.2 (6.4) | 5.8 (5.8) |
| **Refinement** |  |  |  |
| Resolution (Å) | 34.9–1.80 |  | 43.70–2.30 |
| *R*<sub>work</sub>/*R*<sub>free</sub> | 16.7/18.8 |  | 14.3/18.1 |
| **Ramachandran** |  |  |  |
| Favored (%) | 96.7% |  | 96.2% |
| Allowed (%) | 3.3% |  | 3.8% |
| Outliers (%) | 0% |  | 0% |
| **Average B factors (Å<sup>2</sup>)** |  |  |  |
| Protein | 27.2 |  | 30.71 |
| Water | 31.4 |  | 37.43 |
| **RMSDs** |  |  |  |
| Bond lengths (Å) | 0.003 |  | 0.002 |
| Bond angles (°) | 0.562 |  | 0.510 |
| PDB code | 7SB0 |  | 7SB1 |

<sup>a</sup>Values for the highest resolution shells are shown in parentheses.

### The overall structure of acetate-bound CjApe1

CjApe1 has a rigid two-domain structure with an SGNH hydrolase domain (residues S41-Y87 and Y228-Y392) and a CBM35 domain (residues I94-T222) interconnected by two short loops (residues L88-S93 and N223-N227) **(Figure 1)**. An extensive interface, with a buried surface area of 1370 Å^2^ as measured using PISA (28), is found between the SGNH and CBM35 domains. The domain-domain interaction is mediated by a hydrophobic core of 19 hydrophobic amino acids (Ala, Ile, Leu, Met, Val, Trp, Phe, Tyr) from the SGNH domain and 15 hydrophobic amino acids from the CBM35 domain.

The CjApe1 SGNH domain adopts a three-layer α/β/α fold, with a central five-stranded parallel β-sheet (β2, β14, β15, β16, β17) flanked by 10 α-helices (α1-α10) **(Figure 2A, top)**. The invariant catalytic residues (S73, G237, N270 and H369) are situated within a concave surface above the parallel β-sheet of CjApe1 **(Figure 2A, bottom)**. The catalytic triad (S73-D367-H369) form a hydrogen bonding network adjacent to a bound acetate molecule. Clear electron density for the acetate showed that one of the oxygen atoms as 2.7 Å from the amide of G237 and 2.9 Å from Nδ2 of N270, two residues that comprise the oxyanion hole. The methyl group of the acetate packs against a conserved hydrophobic cavity formed by L273 and V368. The orientation of the bound acetate thus likely represents a model for the tetrahedral oxyanion intermediate of the acetyl moiety of the substrate-enzyme complex.

**Figure 2.**
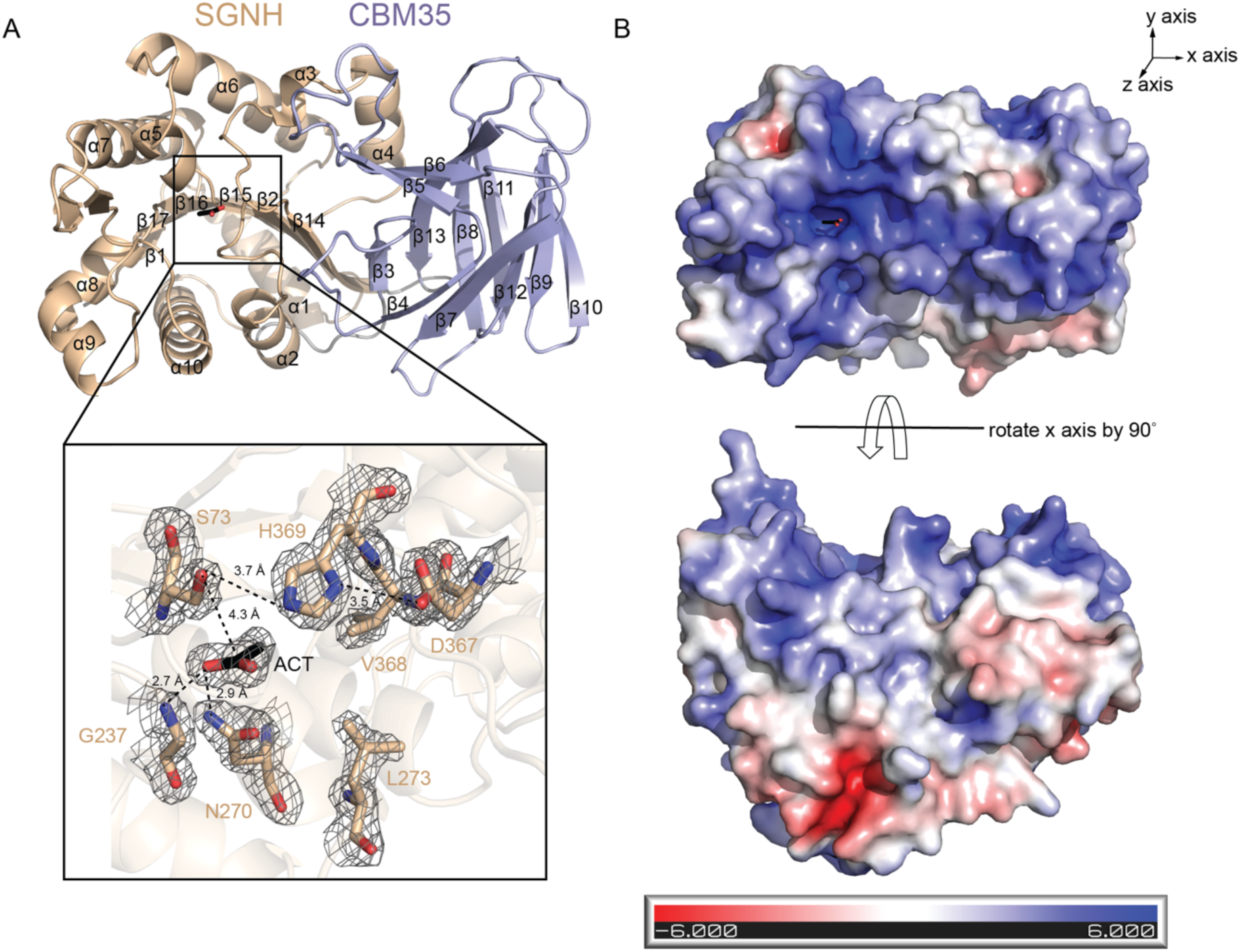
Active site arrangement and electrostatic potential properties of Ape1^41-392^. (A) Magnified view of the active site. The catalytic triad (S73-H369-D367), oxyanion hole (G237 and N270) and conserved hydrophobic residues (V368 and L273) are shown in stick form (nitrogen, blue; oxygen, red; carbon, brown). The electron density is shown as a weighted 2*F*_obs_-*F*_cal_ map contoured at 1 σ. Hydrogen bond networks between residues of the triad are drawn as dashed lines. (B) The electrostatic potential (±6 kT/e) plotted onto the solvent accessible surface of CjApe1. The surface charge was calculated using the APBS plugin in PyMOL, and the input Ape1 structure containing charge and radius information for each atom was prepared using the PDB2PQR web server.

The electrostatic surface potential of CjApe1 was investigated. The side of CjApe with the shallow active site had a largely positively charged surface. Rotating the molecule along the long axis by 90° showed a relatively neutral to negatively charged surface **(Figure 2B)**. The positively charged surface of the CjApe1 may help to orient molecule such that the catalytic cleft is directed towards the negatively charged PG substrate for catalysis.

The CBM35 domain of CjApe1is formed by two sandwiched antiparallel β-sheets (β3, β5, β6, β8, β11, β13 in the first sheet and β4, β7, β9, β10, β12 in the second). A structural homolog search using Dali (29) identified CBM35 domains in various glycoside hydrolases **(Figure 3A–D)**. In these hydrolases, the inter-β-strand loops of the CBM35 domain often coordinate calcium ions and saccharides (26). Saccharide binding by this domain has been proposed to guide substrate specificity of the associated glycoside hydrolase domain (25,30). However, these CBM35 domains and CjApe1 shares sequence identities less than 11% suggesting the CBM35 domain of CjApe1 may have functions other than saccharide binding. Notably, two large loops of the CjApe1 CBM35 domain (CBML1 and CBML2) are situated close to the active cleft of the SGNH domain **(Figure 3E)**. Inspection of the sequences of CBML1 (13 amino acids, A97-N109) and CBML2 (16 amino acids, N121-F136) identified conserved polar (Q105, Q106, N121, S122) and aromatic (Y104 and F132) residues. Inspection of the CjApe1 structure revealed that the side chains of Q105, N121 and R123 form hydrogen bonds with the main chain of the loop in the SGNH domain forming the oxyanion hole (residues A234-D240) **(Figure 3E)**. A similar hydrogen bond network is found in NmApe1 **(Table 2)**. We hypothesize that these residues of the CBM35 domain contributes to enzyme catalysis by stabilizing the oxyanion hole.

**TABLE 2.**
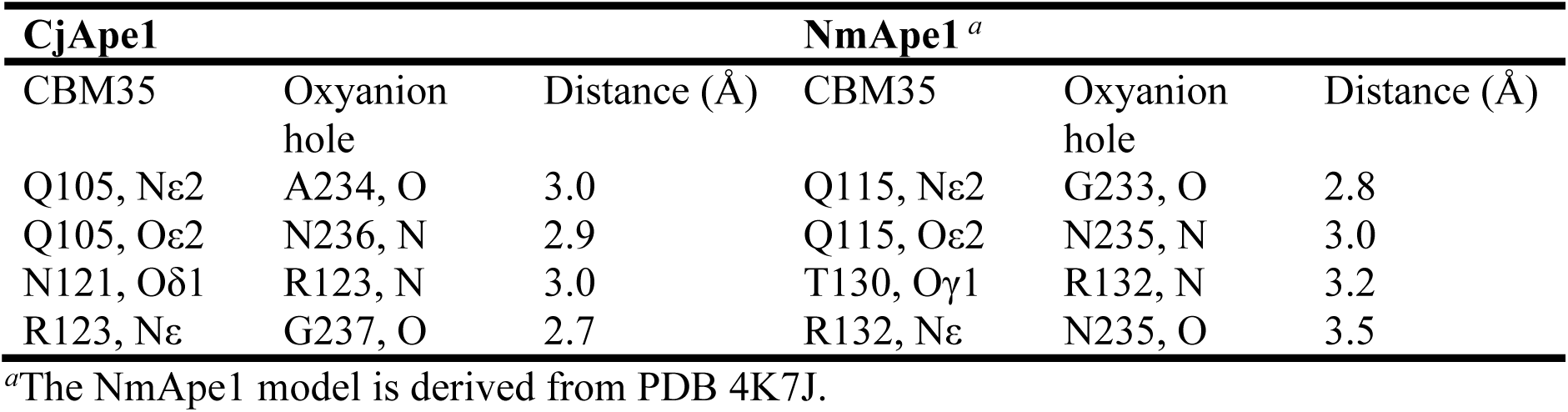
H-bond network between the CBM35 loop and the oxyanion hole

| CjApe1 |  |  | NmApe1 <sup>a</sup> |  |  |
| --- | --- | --- | --- | --- | --- |
| CBM35 | Oxyanion hole | Distance (Å) | CBM35 | Oxyanion hole | Distance (Å) |
| Q105, Nε2 | A234, O | 3.0 | Q115, Nε2 | G233, O | 2.8 |
| Q105, Oε2 | N236, N | 2.9 | Q115, Oε2 | N235, N | 3.0 |
| N121, Oδ1 | R123, N | 3.0 | T130, Oγ1 | R132, N | 3.2 |
| R123, Nε | G237, O | 2.7 | R132, Nε | N235, O | 3.5 |

<sup>a</sup>The NmApe1 model is derived from PDB 4K7J.

**Figure 3.**
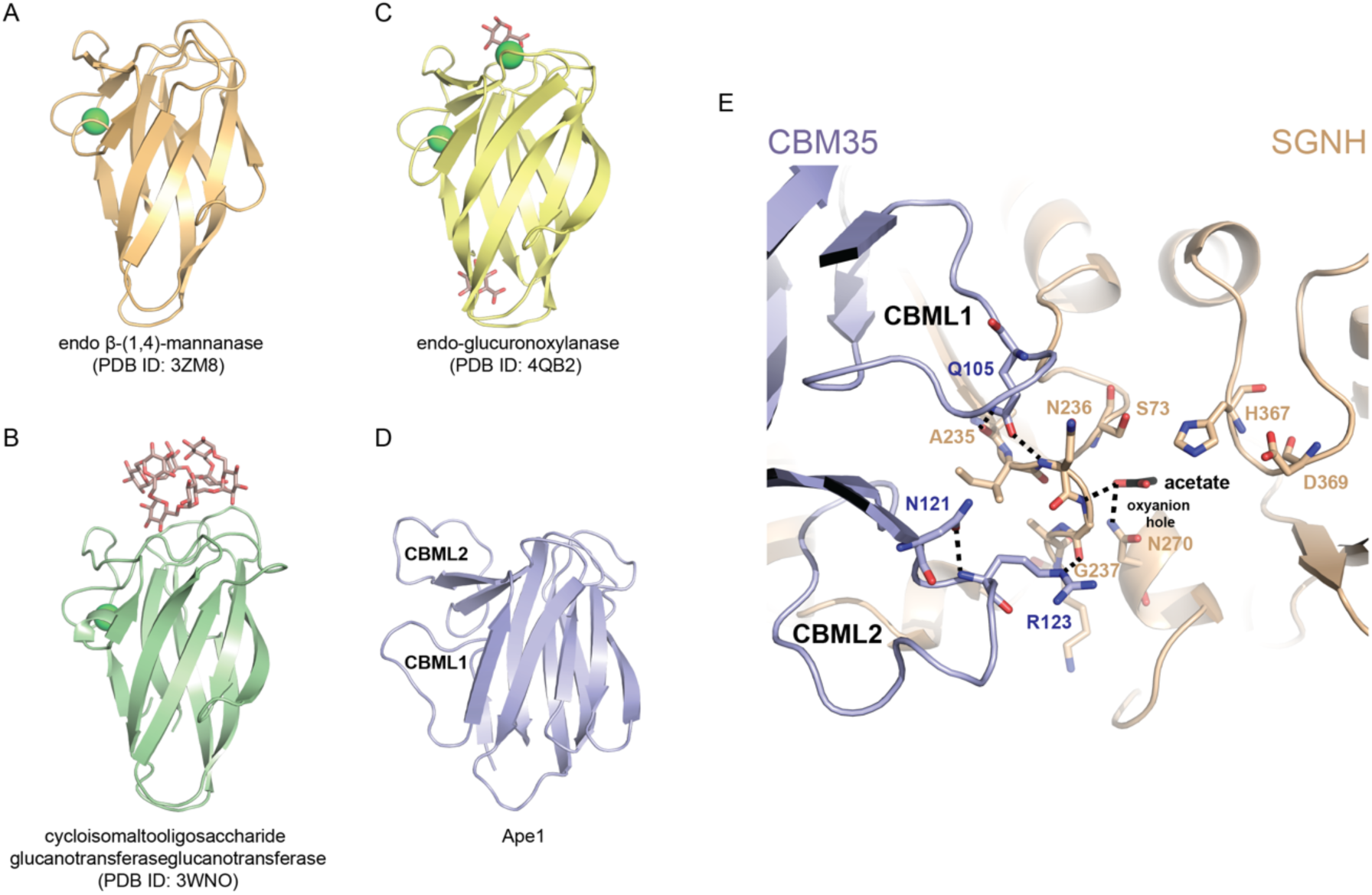
Structural comparison of CBM35 structural homologs. Comparison of CBM35 domain structural homologs: (A) *Podospora anserina*, β-(1,4)-mannanase, PDB 3ZM8; (B) *Paenibacillus barcinonensis*, Xyn30D, PDB 4QB2; (C) *Bacillus circulans*, cycloisomaltooligosaccharide glucanotransferase, PDB: 3WNO; (D) *Campylobacter jejuni*, Ape1. Bound calcium ions and bound saccharides are displayed in green sphere and stick form, respectively. (E) Residue Q105 of CBML1 and residue R123 of CBML2 make H-bonds to residue A235, N236, G237 of the oxyanion hole loop. H-bonds are shown as black dashed lines between atoms.

### Two CBM35 loops promote Ape1 O-acetyl esterase activity

To examine if these loops are required for CjApe1 deacetylase activity, site-directed variants of residues in CBML1 (K103A, Y104G, Q105A, Q106A) and CBML2 (N121A, S122A, R123A, F132A) were generated in the context of the full-length Ape1^22-392^ construct (**Figure 4A)**. All purified CjApe1 variants were concentrated to higher than 9.5 mg/mL, suggesting the proteins are stable in solution. The *O-*acetyl esterase activity of the wild-type and variant proteins were assayed using *p*-nitrophenol acetate (*p*NPAc) as a substrate. Controls with *p*NPAc alone and *p*NPAc incubated with an inactive CjApe1 variant with a substitution of the catalytic nucleophile (S73A) showed minimal absorbance change at 405 nm **(Figure 4B)**. Wild-type CjApe1 cleaved *p*NPAc at a specific activity of 9.6 μmole·min^-1^·mg^-1^ **(Table 3)**. The previously measured CjApe1 activity ranged from 26.1 to 38.9 μmole·min^-1^·mg^-1^ (19) and the activity of NgApe1 is 9.98 μmole·min^-1^·mg^-1^ (14). Wild-type activity levels were observed for the K103A (111%), Y104G (114%), Q106A (103%), S122A (101%) and F132A (96%) variants, indicating that replacement of these residues did not substantially reduce *O*-acetyl esterase activity. On the contrary, reduced deacetylase activity was observed for variants Q105A (59%), N121A (60%), and R123A (34%), suggesting these residues are required for optimal CjApe1 catalysis.

**TABLE 3.**
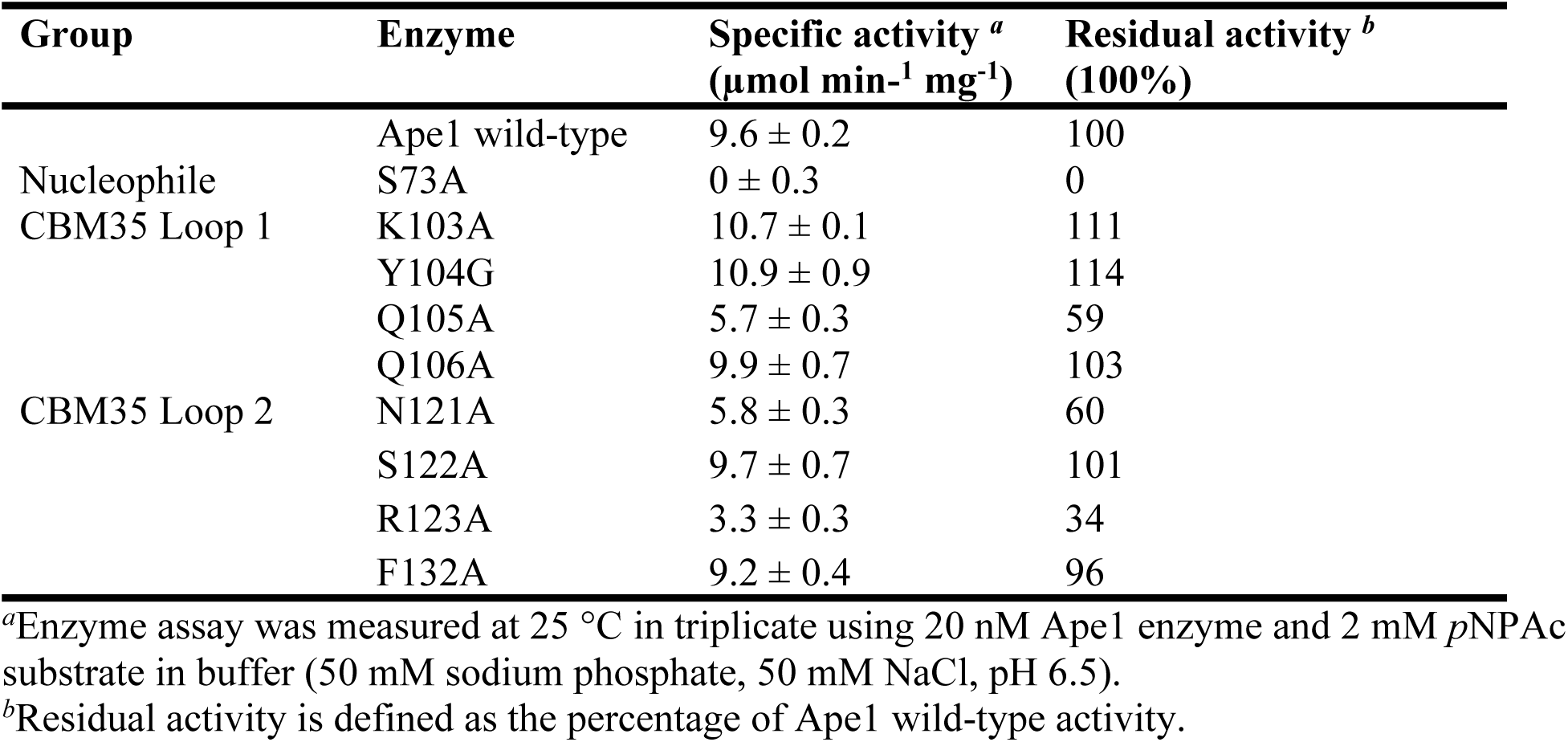
*O*-acetyl esterase activity of CBM35 loop variants on *p*NPAc

**Figure 4.**
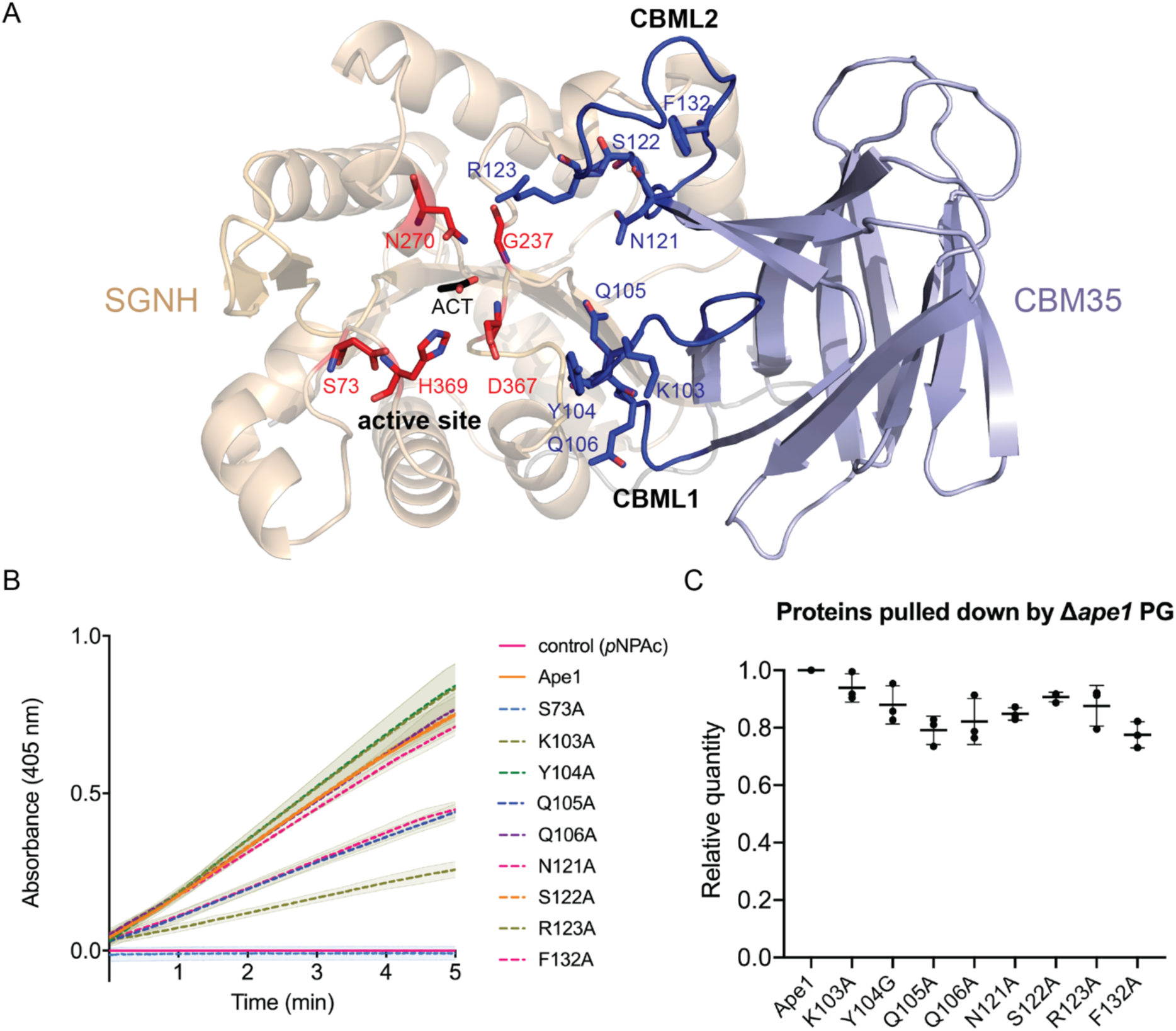
*O*-acetyl esterase activity and PG binding assays of CBM35 loop variants. (A) Residues predicted to be involved in catalysis of *O*-acetylated PG, shown in stick form. The catalytic triad and oxyanion hole are colored in red. CBM35 loops are colored in blue. (B) *O*-acetyl esterase activity of the CjApe1 variants. Purified Ape1 and variants were incubated with *p*NPAc in 50 mM Na_2_HPO_4_/NaH_2_PO_4_, 50 mM NaCl, pH 6.5 at 25 °C for 5 min. The rate of *p*NP generation was monitored spectrophotometrically at 405 nm. Assays were performed in triplicate, mean data are plotted in lines, and errors (within ± standard deviation) are shown as filled area. (C) Binding of CBM35 loop variants to Δ*ape1* PG. Wild-type and variant proteins were incubated with *C. jejuni* Δ*ape1* PG in 50 mM Na_2_HPO_4_/NaH_2_PO_4_, 50 mM NaCl, pH 6.5 at 4 °C for 20 min. Unbound proteins were washed with buffer and removed by centrifugation. Proteins pulled-down by PG were recovered using buffer and analyzed by SDS-PAGE. Band intensities were quantified using ImageJ. The relative quantity (%) was calculated as amounts of pulled-down variants relative to pulled-down wild-type (set as 1). Assays were performed in triplicate, and data are shown as mean ± the standard deviation. Each dot reflects an individual experiment.

To examine whether the reduced activity of the variants Q105A, N121A, and R123A was due to a deficiency in PG-binding, a PG pulled-down assay was performed. CjApe1 and variant proteins were incubated with insoluble PG isolated from *C. jejuni* Δ*ape1*, followed by centrifugation and wash steps. Proteins pulled down by Δ*ape1* PG were recovered from the pellet and quantified using SDS-PAGE. Protein pulled down in the absence of PG was minimal **(Figure S1A)**. Conversely, CjApe1 and variants were pulled down by Δ*ape1* PG. Relative PG binding was estimated by band densitometry analysis by comparing the amount of variant protein pulled down to that of wild-type CjApe1 **(Figure S1B)**. Slight reductions were observed for all variants tested: K103A (94%), Y104G (88%), Q105A (90%), Q106A (82%), N121A (85%), S122A (91%), R123A (88%) and F132A (78%) **(Figure 4C)**; however, the discernable differences in the amount of protein pulled down did not explain the larger differences in *p*NPAc activity. The site-directed variants support a role for the CBML1 and CBML2 loops in deacetylase catalysis.

### Ape1-PG binding mode by mechanism-guided HADDOCK docking

To obtain a plausible binding mode of CjApe1 to PG, docking experiments were performed using HADDOCK2.2 with active site restraints (31,32). The CjApe1 crystal structure and an ensemble of 10 *O*-acetyl PG conformers were used for docking. The *O*-acetyl PG model was prepared as a hexasaccharide to approximately match the length of the putative substrate binding groove (20-30Å) on the CjApe1 protein surface. Considering ∼10% of total Δ*ape1* PG is acetylated (19), only the second MurNAc residue in the hexasaccharide was acetylated. A total of six unambiguous distance restraints were applied for the CjApe1-PG complex to maintain catalytically reasonable distances between the catalytic triad, oxyanion hole and the acetyl group of the hexasaccharide (see methods for details).

Of the final 200 docking solutions, 188 were grouped into 7 clusters using a fraction of common contacts (FCC) cut-off of 0.79 **(Table S2)**. All of the clusters showed a convergent binding mode with an RMSD < 2.5 Å to the model with the best HADDOCK score (lowest energy) **(Figure S2A)**. Importantly, each cluster featured the *O*-acetyl glycan strand sitting along the interface of the SGNH and CBM35 domains **(Figure 5)**. The main difference among the 7 clusters was the direction of *O*-acetyl hexasaccharide **(Figure S2B)**, in which the reducing end of the hexasaccharide, containing the free hydroxyl group at carbon 1, is either close to the CBML1 loop (named as O1→O4) or close to the CBML2 loop (named as O4←O1) **(Figure 5)**. 91% of the 188 solutions, including the best HADDOCK score, belongs to O1→O4; 9% of the 188 solutions were oriented as O4←O1. However, the direction preference cannot be discerned in the docking experiment. Repeating the docking experiment with another random set of simulated PG conformers resulted in the highest scoring model with the opposite glycan direction, suggesting the glycan orientation bound to Ape1 varies depending on the input glycan conformers. Between the top docking solutions, only subtle changes were observed in the phi-psi angle of glycan strands. The phi-psi angle of the bound glycan strand is comparable to that of the starting backbone of glycan strand conformers, suggesting the binding pose maintains the low energy conformer state of the PG hexasaccharide.

**Figure 5.**
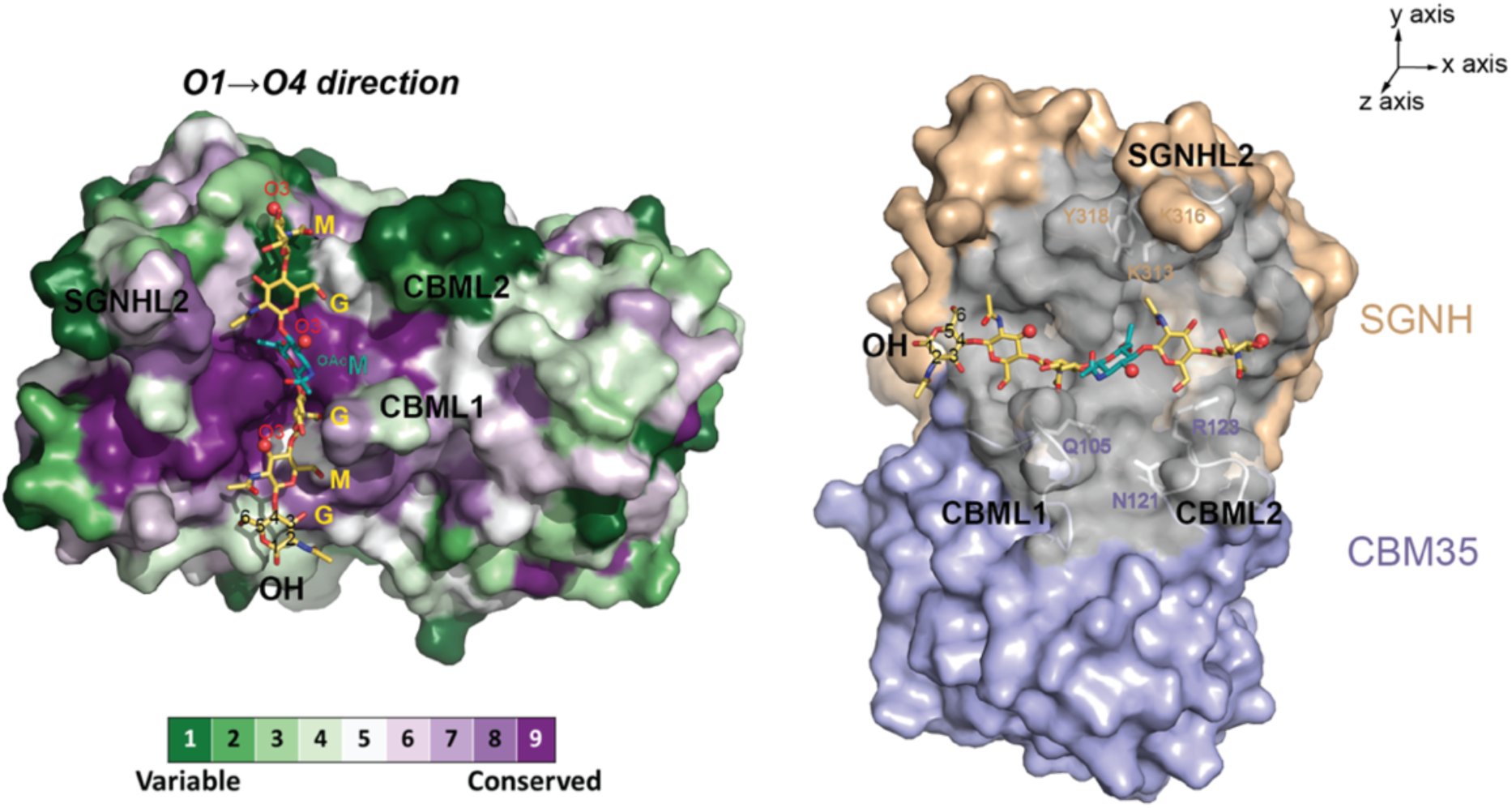
HADDOCK model of the CjApe1-*O*-acetyl hexasaccharide complex. The CjApe1-*O*-acetyl hexasaccharide model presents the best HADDOCK scoring solution in the docking experiment. Two views are rotated by 90° along the z axis as to emphasize that the proximal loops form a clamp that holds a hexasaccharide. In the left panel, CjApe1 is shown in the surface representation colored by amino acid conservation. The *O*-acetyl hexasaccharide is shown in stick form and saccharide residues (MurNAc=M; GlcNAc=G; *O*-acetylated MurNAc=OAcM) are labelled. MurNAc O3 atoms are highlighted as red spheres. In the right panel, residues that are within 15 Å to *O*-acetylated MurNAc are colored in grey, including SGNHL1, CBML1 and CBML2. The predicated functional residues in these loops are labelled.

The protein surface of a representative complex **(Figure 5, left)** is colored by the amino acid conservation derived from ConSurf analysis (33). The bound glycan strand runs the length of the putative substrate groove on the CjApe1 surface, and the *O*-acetylated MurNAc sits adjacent to the catalytic center of the enzyme. Each MurNAc O3 atom is exposed to the solvent, allowing the peptide stem to point away from Ape1 without steric clashes. In the docked model, surface loops of CjApe1 are predicted to act like clamps that hold the glycan strand in place adjacent to the catalytic residues. Loops with residues less than 15 Å from the bound *O*-acetylated MurNAc include SGNHL1 (loop that connects β2 and α2), SGNHL2 loop (between β16 and α7) and the two CBM35 loops CBML1 and CBML2 **(Figure 5, right)**. Loop SGNHL2 contains several conserved residues (K313, Y316 and K318), which may be involved in Ape1 activity through an unidentified mechanism. Residues Q105, N121 and R123 of CBML1 and CBML2, in which mutations reduced catalytic activity, are within 15 Å from *O*-acetylated MurNAc.

### Structural comparison of Ape1 homologs

To study whether the loops in the active site of CjApe1 determine substrate entry and binding, CjApe1 was compared to its structural homologs. From a search in the PDB database using Dali, the NmApe1 (PDB ID: 4K7J) is the closest structural homolog of CjApe1, with an RMSD of 2.0 Å over 267 aligned Cα positions. A carbohydrate esterase of the SGNH superfamily from *Clostridium thermocellum* (CtCE2; PDB ID: 2WAB) showed an RMSD of 2.7 Å over 98 aligned Cα positions.

The concave surface that forms the active groove of the CjApe1 and NmApe1 are generally similar **(Figure 6A-B)**. Both CjApe1 and NmApe1 contain long SGNHL2 (17 amino acids) and short SGNHL1 loops (8 amino acids). CjApe1 and NmApe1 had the largest structural deviation at loop SGNHL2. In NmApe1, the SGNHL2 loop conformation is nearly 90 ° from of the equivalent position of this loop in CjApe1 **(Figure 7A-B)**. The difference in the conformation of this loop is likely caused by a disulfide bond between Cys316 and Cys352 observed in the NmApe1 structure (PDB ID: 4K7J). The cysteine residues are conserved in several bacteria but not in *C. jejuni* **(Figure 7C and Figure S3)**.

**Figure 6.**
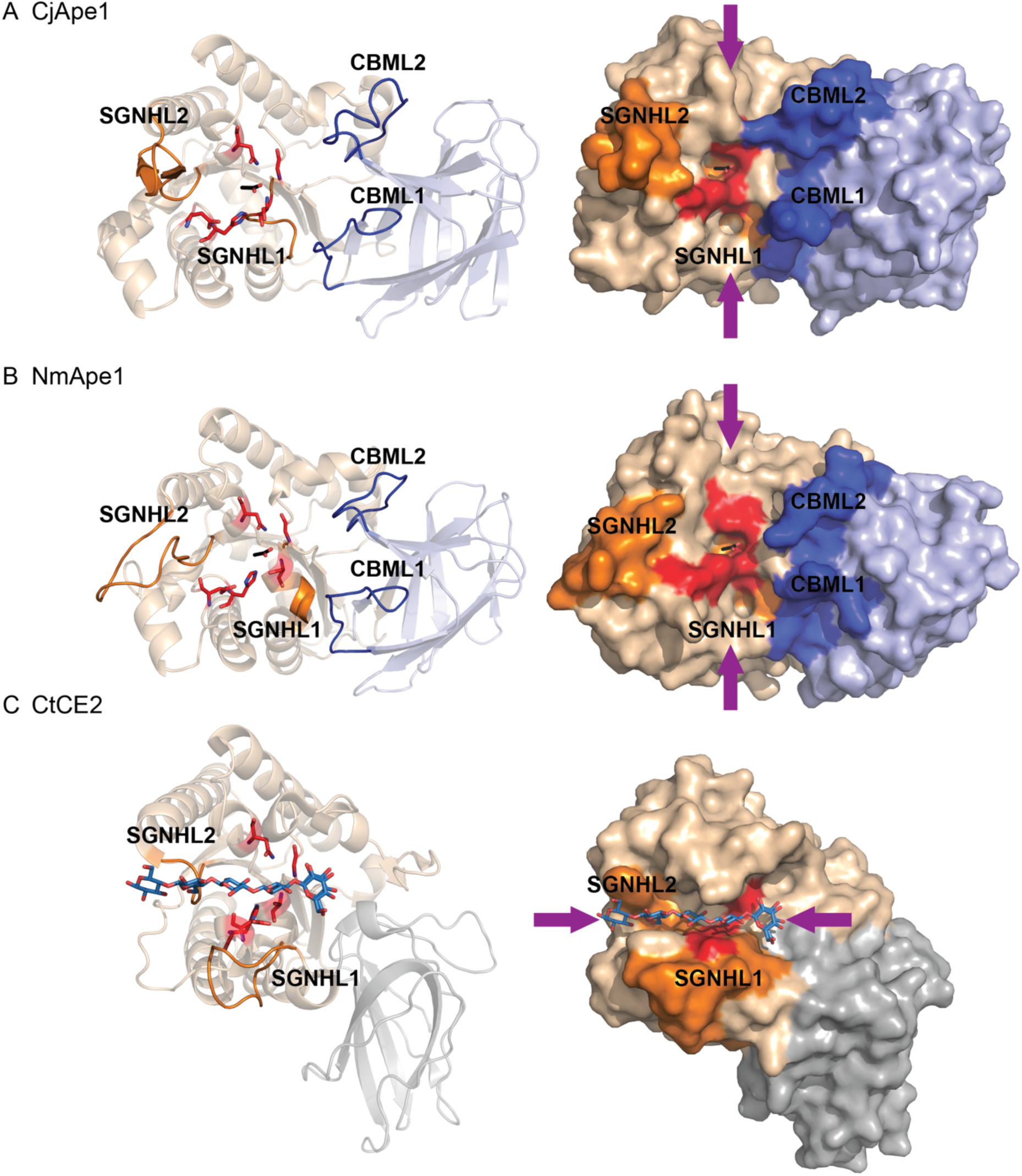
Comparison of proposed substrate binding orientations within the SGNH hydrolase superfamily. Structures of (A) CjApe1, (B) NmApe1 (PDB 4K7J) and (C) CtCE2 (PBD: 2WAB) are shown as cartoons (left) and surface representations (right). These structures are aligned based on the orientation of the SGNH domains. The orientation of glycan substrates bound in the active site grooves are indicated using purple arrows. Loops involved in determining the substrate orientation are highlighted (SGNHL1 and SGNHL2, orange; CBML1 and CBML2; blue). Conserved SGNH domain catalytic residues are shown in stick form and colored red.

**Figure 7.**
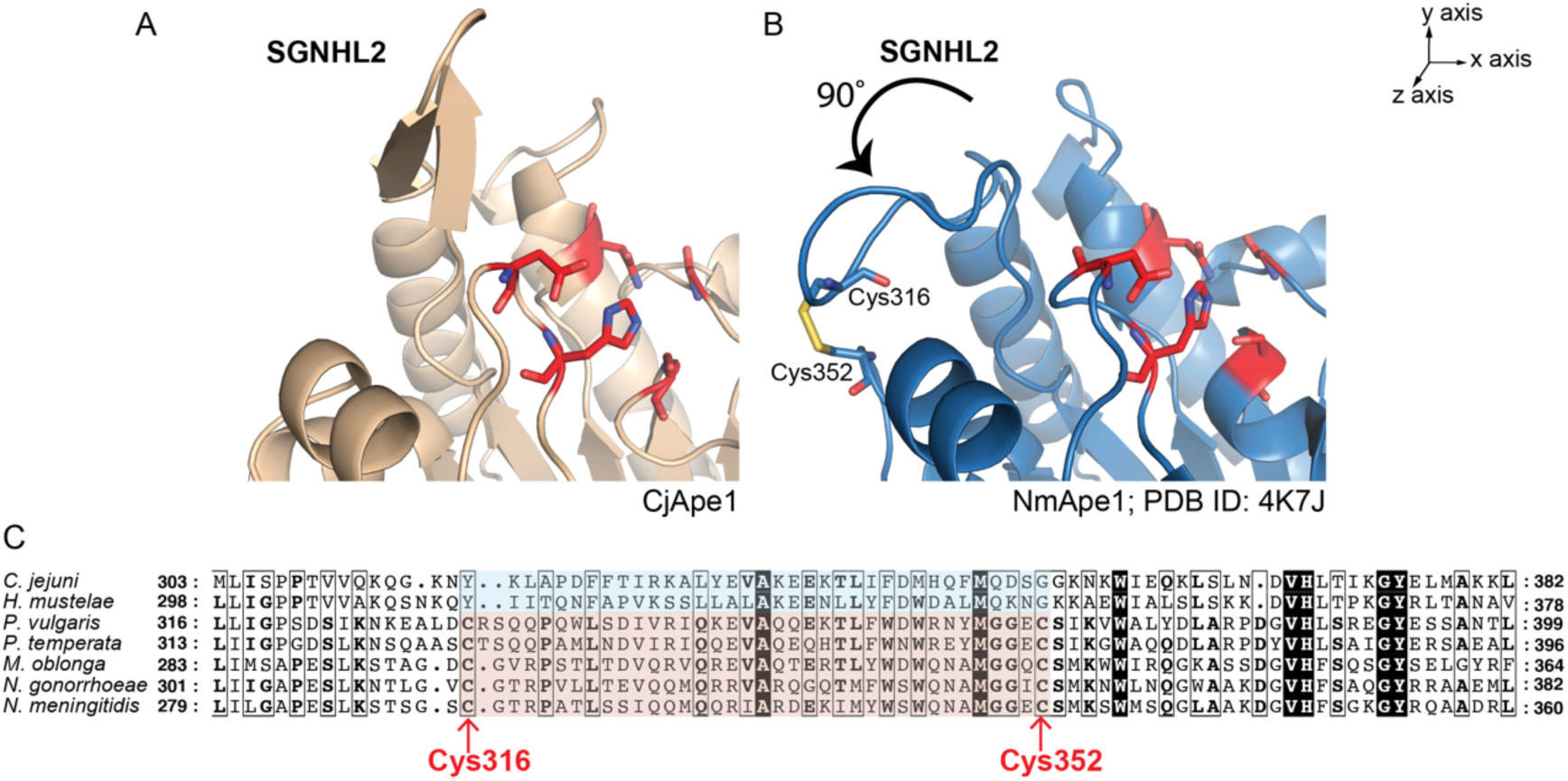
Structure of the SNGHL2 loop and sequence alignment. (A) Cartoon representation of the SGNHL2 loop from CjApe1. Conserved SGNH domain catalytic residues are shown as sticks and colored red. (B) NmApe1 (PDB 4K7J blue; 4K9S green). The SGNHL2 loop of CjApe1 is orientated 90° about the z-axis in comparison to the loops in the NmApe1 structure. Residues constituting the disulfide bond are shown in stick form (nitrogen, blue; oxygen, red; sulfur, yellow). (C) Sequence alignment of CjApe1 homologs performed in Clustal Omega (45) and presented by ESPript (46). Highlighted in this partial alignment is the Cys-Cys bond of the SGNHL2 loop. Ape1 homologs: *Helicobacter mustelae* (WP_013022746.1); *Proteus vulgaris* (WP_185901317.1); *Photorhabdus temperata* (WP_023045068.1); *Moraxella oblonga* (WP_066805747.1); *Neisseria gonorrhoeae* (WP_003695451.1); *Neisseria meningitidis* (WP_049224741.1).

The surface of CtCE2 near the active site is significantly different as compared to CjApe1 and NmApe1. In CtCE2, the SGNHL2 loop is 6 residues long, which is 11 amino acids shorter than in CjApe1. Conversely, CtCE2 loop SGNHL1 is 4 amino acids longer than the equivalent loop in CjApe1 **(Figure 6A & 6C)** The shorter SGNHL2 and longer SGNHL1 loops of CtCE2 restructure the substrate binding groove to be 90° from that in Ape1. The consequence of the altered loops is illustrated by the orientation of bound cellulohexose to CtCE2 which is perpendicular in comparison to the glycan in the CjApe1 docking model **(Figure 6C)**. Collectively, we propose the orientation of substrate binding groove in Ape1 is divergent from that of CtCE2 because of altered loop lengths in the active cleft. Such changes might indicate an adaptation to substrate specificity.

## Discussion

CBM35 domains are proposed to bind sugars and assist in catalytic efficiency of glycoside hydrolases (26,30). The domain displays a jelly roll/β-sandwich fold with two conserved calcium ions binding sites. The first calcium ion is considered a structural site coordinated by conserved acidic residues. The second calcium ion is typically involved in sugar binding. The bound sugar is coordinated by the calcium ion and interacts with conserved residues such as stacking interactions with aromatic residues (Trp and Phe) (25,30,34) and charged interactions with arginine residues (25,26). The structure of CBM35 in CjApe1 diverges from CBM35 domains in other glycoside hydrolases. We did not observe bound metal ions nor the conserved amino acids for metal ion binding, consistent with the observation that EDTA treatment of NgApe1 did not inhibit Ape1 activity (21). The absence of metal binding is reflected in the low level of sequence identity of the Ape1 CBM35 and the CBM35 domains from other glycoside hydrolases.

We showed that the CjApe1 CBM35 has two big loops positioned near to the catalytic triad. The length of these loops is longer than the equivalent loops in other CBM35 domains suggesting that these long loops evolved for distinct biological function. The catalytic domain of SGNH hydrolase superfamily is characterized with a canonical α/β/α fold. Interestingly, The CjApe1 CBM35 domain is inserted between helix α2 and strand β14 in the α/β/α fold. This insertion places the big loops of the CBM35 domain in proximity to the active site. Our mutagenesis results confirmed that residues of the CjApe1 CBM35 domain are important for catalysis. The insertion of CBM35 domain might be an adaptation for CjApe1 for hydrolysis of the PG substrate.

The proposed CjApe1-PG binding model features a long putative substrate binding groove docked with a six-saccharide polymer. *C. jejuni* Δ*ape1* showed increased *O*-acetylated of MurNAc linked to dipeptides, tripeptides, and tetrapeptides when compared to the wild-type strain (19), suggesting that muropeptide length has little effect on CjApe1 de-*O*-acetylase activity. Purified NmApe1 is active against various *O*-acetylated muropeptides *in vitro*, but deletion of *N. meningitidis ape1* displayed accumulated *O*-acetylated tripeptide levels, suggesting a preference for *O*-acetylated tripeptides in cell (20). However, the mechanism leading to a preference for tripeptide substrates in *N. meningitidis* is not known. Our Ape1-PG complex model suggests that the peptides are positioned away from the protein and do not form specific interactions. Ape1 is proposed to act as a prerequisite enzyme for LT to control glycan strand length in the cell (15,19,20). LTs cleave the glycosidic bond between MurNAc and GlcNAc, and catalyze the concomitant formation of a 1,6-anhydroMurNAc end. During PG turnover, CjApe1 might efficiently de-*O*-acetylate PG containing glycan strands that are longer than six saccharides and initiate LT activity for subsequent biological events.

A direct binding between Ape1 and LT was recently identified by gel filtration in *N. meningitidis*, revealing a 105 kDa complex from co-elution of NmApe1(40 kDa) and the LT LtgA (65 kDa) (35). The authors monitored NmApe1 *O*-acetyl esterase activity on *p*NPAc in the presence of LtgA, finding that maximal NmApe1 activity is dependent on the presence of LtgA. BLAST searches with LtgA in *C. jejuni* found Slt (*CJJ81176_0859*) shares 30% with LtgA, and the recombinant Slt was expressed as described (36). We did not observe a change in CjApe1 *O*-acetyl esterase activity in the presence of Slt **(Figure S4)**. This suggests that the mechanisms of Ape1 in *C. jejuni* and *N. meningitidis* are distinct, possibly due to requirements of helical shape maintenance in *C. jejuni* that are absent in spherical *N. meningitidis*.

The SGNHL2 loop of CjApe1 displayed distinct conformation from that of NmApe1**(Figure 7)**. It is important to note that this disulfide bond in SGNHL2 is conserved among NmApe1 homologs from betaproteobacteria and gammaproteobacteria but is absent in epsilonproteobacteria **(Figure S3)**. In the CjApe1 and in other *Campylobacter* and *Helicobacter* species, both Cys residues are absent from the SGNHL2 loop. In NgApe1, titration of 5,5’-dithiobis-(2-nitrobenzoic acid) to quantify free thiolate groups indicated that ∼65% of SGNHL2 Cys residues formed a disulfide bond (37). The dynamics of the Cys redox state of NgApe1 (and NmApe1) is hypothesized to regulate its activity. Upon the treatment of NgApe1 with thiol oxidizing reagent diamide, activity on *p*NPAc was reduced by 70%. Treatment of NgApe1 with the reducing agent glutathione also showed a 30% reduction in activity. Future work on exploring the function of the SGNHL2 loop on Ape1 activity on PG substrates would help to understand possible regulatory mechanisms of Ape1 catalysis. Key residues to study by site-directed mutagenesis include introducing Cys residues at equivalent positions in CjApe1 and the substitutions at conserved positively charged residues of the SGNHL2 loop **(Figure 5, right)**.

In our CjApe1-PG model, the binding interface consists of a positively charged groove in CjApe1 and the glycan saccharides of PG. A high throughput inhibitor screen using fluorogenic substrate 4-methyllumbelliferyl acetate (MU-Ac) identified 7 potential NgApe1 inhibitors (37). These compounds feature phenyl rings and hydroxyl groups, which show similarity to the saccharide structure of PG. One of the inhibitors is purpurin (*K*_i_=4.8 μM), an anthraquinone-based compound found in the roots of the plant *Rubia tinctorum*. Attempts to obtain a structure of the CjApe1-purpurin complex was performed by soaking and co-crystallization experiments.

Crystals that were soaked with purpurin turned from clear to yellow in color, suggesting possible binding. Crystals of Ape1 were obtained from solution in the presence of inhibitor. However, these crystals showed poor diffraction to ∼6.5 Å, possibly due to local conformational changes disrupting crystal packing upon purpurin binding.

## Experimental procedures

### Cloning

A list of strains and primers used in this study can be found in **Table S1**. The expression vector pET28a-Ape1^22-392^ was donated by Dr. Erin Gaynor (19). The encoded product includes an N-terminal poly-His tag followed by a thrombin cleavage site and the full-length Ape1 protein without the N-terminal signal peptide (residue 1-21). The expression vector pET28a-Ape1^41-392^ encoding Ape1 protein with an N-terminal His_6_-tag followed by a thrombin cleavage site and residues 41–392 was constructed using the restriction enzyme double-digestion method. The portion of *ape1* (*CJJ81176*_0638) corresponding to the product without the N-terminal signal peptide and the subsequent 19 residues was amplified from *C. jejuni* 81-176 genomic DNA using primers Ape1^41-392^(F) and Ape1^41-392^(R). The PCR product and pET-28a(+) vector were digested with restriction enzymes NheI/BamHI, and ligated together by T4 DNA ligase (NEB). The recombinant plasmid was then transformed into *E. coli* DH5α, selected using kanamycin, and validated by PCR analysis and sequencing.

Site-directed mutagenesis was used to generate the CBM35 loop variants K103A, Y104G, Q105A, Q106A, N121A, S122A, R123A, F132A using pET28a-Ape1^22-392^ as a template. Each primer was 5′ phosphorylated and designed to contain a complementary mutation of the target sequence. Whole-plasmid amplification reactions were performed using Phusion polymerase (NEB) and Ampligase (Epicentre). Reactions were digested with DpnI for 3 hours at 37 °C to remove methylated template vectors.

### Recombinant protein expression and purification

Ape1^22-392^ and Ape1^41-392^ proteins were prepared in *E. coli* BL21(DE3) grown overnight at 37 °C in Luria Bertani (LB) media containing 25 μg/ml kanamycin. Overnight cultures were inoculated into 1 L LB media at a 1:100 dilution and grown at 37 °C to OD_600_ of 0.8-1.0 before induction with 0.5 mM isopropyl β-D-thiogalactopyranoside (IPTG) at 20 °C for 16 hours. The cells were pelleted by centrifugation at 5,000 rpm at 4 °C for 15 min. The cell pellet was resuspended at 4 °C in 50 mM Tris-HCl pH 7.0, 500 mM NaCl, 20 mM Imidazole, 1 mM phenylmethylsulfonyl fluoride (PMSF), and DNase and lysed using an Emulsi Flex-C5 homogenizer (Avestin). The cell lysate was centrifuged at 16,000 rpm at 4 °C for 50 min, then the soluble fraction was filtered through 0.22 μm PVDF membrane before loading onto a 5 mL HisTrap HP column (GE Healthcare). The column was washed with 20 column volumes of 50 mM Tris-HCl pH 7.0, 500 mM NaCl, and Ape1 was eluted with imidazole. Ape1 was dialyzed against 20 mM Tris-HCl pH 7.0, 150 mM NaCl and simultaneously digested with thrombin (200:1 w/w Ape1: thrombin ratio) at 4 °C overnight to remove the His_6_-tag. The Ape1-thrombin mixture was incubated with p-aminobenzamidine-agarose beads (5 mg thrombin:1 mL beads ratio; Sigma) at 4 °C for 15 min to remove thrombin and was then filtered through a 0.22 μm PVDF membrane. The cleaved protein was separated using a second HisTrap HP column (GE Healthcare). His_6_-tag free Ape1 protein was then loaded onto a Superdex 200 10/300 GL column (GE Healthcare) in 20 mM Tris-HCl pH 7.0, 150 mM NaCl. Finally, monodisperse tag-free Ape1 was concentrated using 10 kDa MWCO Amicon (Millipore) to 15–20 mg/ml, flash frozen in liquid nitrogen, and stored in −80 °C. Protein purity was assessed by SDS-PAGE and electrospray ionization mass spectroscopy (MSL/LMB Proteomics Core Facility, UBC).

SeMetApe1^41-392^ was prepared in *E. coli* BL21(DE3) grown overnight at 37 °C in LB media containing 25 μg/ml kanamycin. Pelleted cultures were then inoculated in 1 L M9 minimal media (6 g Na_2_HPO_4_, 3 g KH_2_PO_4_, 1 g NH_4_Cl, 0.5 g NaCl, 1 mM MgSO_4_, 40% (w/v) glucose, 0.5% (w/v) thiamine, 4.2 g Fe_2_SO_4_, 25 μg/mL kanamycin per liter) and grown at 37 °C to an OD_600_ of 0.3. 100 mg of L-lysine, L-threonine, L-phenylalanine and 50 mg of L-isoleucine, L-leucine, L-valine, and L-seleno-methionine were then supplemented into the 1 L culture media, followed by induction with 0.5 mM IPTG at 20 °C for 16 hours. SeMetApe1^41-392^ was purified using the same protocol as for unlabeled protein.

### Crystallization and Structure determination

Ape1^41-392^ was crystallized by hanging drop vapor diffusion. The crystallization well contained a 900 μl solution of 100 mM CAPS pH 10.5, 200 mM NaCl, 16% (w/v) PEG8000, 2.5% (w/v) PEG3350. 1 μL of this solution was mixed with 1 μL Ape1^41-392^ (12 mg/ml). A rod-shaped crystal with a size of 0.2 μm appeared after one day of incubation at room temperature. The crystal was submerged in 35% (w/v) PEG8000 prepared in the crystallization solution as a cryoprotectant, then immediately stored in liquid nitrogen before data collection. A native dataset was collected at 1 Å at 100 K on beamline 9-2 at the Stanford Synchrotron Radiation Lightsource (SSRL; Palo Alto, CA). SeMetApe1^41-392^ was crystallized as described for unlabeled Ape1^41-392^ with modification. The crystallization well contained 100 mM CAPS pH 10.5, 200 mM NaCl, 16% (w/v) PEG 8000, 12 mM phenol. A 0.4 mm rod-shaped crystal was submersed in cryoprotectant consisting of 35% (w/v) PEG 8000 and 10% (v/v) glycerol prior to freezing. A single-wavelength anomalous dispersion (SAD) dataset was collected at 0.979 Å.

The collected datasets were indexed, integrated, and scaled with HKL2000 (38). Structural determination was conducted using software packages in Phenix (39). SeMetApe1^41- 392^ SAD dataset was processed in phenix.AutoSol to obtain initial phases and a preliminary model through automated fitting. Manual model construction of SeMetApe1^41-392^ was done with Coot (40) and phenix.refine. The Ape1^41-392^ native dataset was phased by molecular replacement, and the model was built as described above. The side chains of K49, Q63, K224, K316 of chain A to C; K177, K313, Q314 and K319 of chain A; K147 and K313 of chain B; K50, K53, E56, K168, K275, K298, Q361 and D391 of chain C were not modelled due to disorder.

Ape1^22-392^ was crystallized using hanging drop vapor diffusion. The crystallization well contained 900 μl of 166 mM sodium acetate, 28% (w/v) PEG4000, and 80 mM Tris-HCl pH 8.5. 1 μl precipitant solution and 1 μl Ape1^22-392^ (28 mg/mL) were mixed with 0.2 μl of 10 mM GSH (L-Glutathione reduced) and 10 mM GSSG (L-Glutathione oxidized) in the hanging drop. Crystals grew at room temperature within a week. The crystal was briefly soaked in crystallization solution containing 15% (v/v) glycerol before stored in liquid nitrogen. A native dataset was collected at 0.98 Å at 100 K at the Canadian Light Source on beamline 08ID-1. Data process, phasing and model building are same to methods of the Ape1^41-392^ model determination. Side chains of K329 of chain B and D28 of chain A to C were not included in the final model due to disorder.

### O-acetyl esterase activity on 4-Nitrophenyl acetate

*O*-acetylesterase activity was quantified using a colorimetric assay. 20 nM Ape1 protein and 2 mM *p*NPAc (prepared in methanol and diluted with reaction buffer to a final concentration of < 1%) were incubated in a 300 μl volume of 50 mM sodium phosphate pH 6.5 and 50 mM NaCl at 25 °C for 5 min. 1 unit of specific activity was defined as the amounts of released *p*-nitrophenol (μmole) per min per mg of protein. A molar extinction coefficient of 18,000 M^-1^cm^-1^ for *p*-nitrophenol at 405 nm was used to calculate product formation (41).

### Peptidoglycan pull-down

40 μg of Ape1 or variant proteins were incubated with 50 μg of purified *C. jejuni* Δ*ape1* PG in 250 μL of reaction buffer (50 mM sodium phosphate pH 6.5, 50 mM NaCl) at 4 °C for 20 min, followed by centrifugation at 13,000 rpm for 10 min. To remove unbound proteins, insoluble PG and pulled-down proteins were washed three times with 1 mL of buffer. Pulled-down proteins were analyzed by SDS-PAGE. Band intensities were quantified in ImageJ.

### HADDOCK docking

A model of the CjApe1-PG complex was produced using HADDOCK 2.2 (31,32). The Ape1 docking conformer was extracted from the crystal structure of the acetate-bound CjApe1^41- 392^, with missing side-chains rebuilt using Coot (40). The binding interface of Ape1 was predicted using the CPORT server (42). CPORT predicts the consensus binding interface from the results of 6 prediction servers. An output of active (i.e., involved in binding) and passive residues (∼5 Å proximal to the binding site) were produced from CPORT.

An ensemble of 10 *O*-acetyl hexasaccharide conformers were used to represent the PG glycan. The phi/psi angles of the β-1,4 glycosidic bond in the hexasaccharide was modelled at 69°/12°, as was previously determined by NMR (43). An *O*-acetyl group was manually built onto the second MurNAc residue of the hexasaccharide using CNS (44). The ensemble of *O*-acetyl hexasaccharide conformers were generated by simulated annealing and energy minimization in CNS. All residues in *O*-acetyl hexasaccharide were defined as passive and are fully flexible in HADDOCK.

The list of Ambiguous interaction residues (AIR) is summarized in **Table S2**. A total of 6 unambiguous distance restraints were used in docking. Two involved the triad hydrogen bond distance: 2.5–3.5Å between Oδ2 of the acid D367 and Nδ1 of the base H369, and 3.5 Å between Nε2 of the base H369 and Oγ of the nucleophile S73. The remaining four involved the bond distance between the catalytic cleft and the *O*-acetyl group of the substrate, including 2.5–3.5Å between the Oγ of S73 and carbonyl carbon of *O*-acetate, 2.5 Å between the *O*-acetate oxygen atom to the amide nitrogen of G237 and Nδ2 of N270 from the oxyanion hole, and 4.0 Å of hydrophobic contact between the methyl group carbon of *O*-acetate and Cγ of V368. In the docking procedure, a sample of 10,000 docking solutions were generated at the rigid body stage. The top 400 complexes based on HADDOCK scoring were subjected to simulated annealing and the resulting top 200 complexes were further refined with waters.

## Acknowledgements

Use of the Stanford Synchrotron Radiation Lightsource, SLAC National Accelerator Laboratory, is supported by the U.S. Department of Energy, Office of Science, Office of Basic Energy Sciences under Contract No. DE-AC02-76SF00515. The SSRL Structural Molecular Biology Program is supported by the DOE Office of Biological and Environmental Research, and by the National Institutes of Health, National Institute of General Medical Sciences (P41GM103393).

The contents of this publication are solely the responsibility of the authors and do not necessarily represent the official views of NIGMS or NIH. Research described in this paper was performed using Beamline 08B1-1 at the Canadian Light Source, a national research facility of the University of Saskatchewan, which is supported by the Canada Foundation for Innovation (CFI), the Natural Sciences and Engineering Research Council (NSERC), the National Research Council (NRC), the Canadian Institutes of Health Research (CIHR), the Government of Saskatchewan, and the University of Saskatchewan.

## Funding and additional information

This research was funded by Canadian Institutes of Health Research (CIHR) grants MOP-142176 to M.E.P.M. Instrument support was provided by the Natural Sciences and Engineering Research Council (NSERC) of Canada and the University of British Columbia. The funders had no role in study design, data collection and interpretation, or the decision to submit the work for publication.

