## Supporting information for "Mechanism of the CBM35 domain in assisting catalysis by Ape1, a *Campylobacter jejuni* O-acetyl esterase"

**Running title:** CBM35 assists Ape1 activity

Supporting information

Table S1. Strains and primers used in this study

| **Strain** | **Description** |
| --- | --- |
| BL21-Ape1^22-392^ | *E. coli* BL21(DE3) strain harboring plasmid pET28a-Ape1^22-392^ *^a^* |
| BL21-Ape1^41-392^ | *E. coli* BL21(DE3) strain harboring plasmid pET28a-Ape1^41-392^ |
| BL21-Ape1^K103A^ | *E. coli* BL21(DE3) strain harboring plasmid pET28a-Ape1^K103A^ |
| BL21-Ape1^Y104G^ | *E. coli* BL21(DE3) strain harboring plasmid pET28a-Ape1^Y104G^ |
| BL21-Ape1^Q105A^ | *E. coli* BL21(DE3) strain harboring plasmid pET28a-Ape1^Q105A^ |
| BL21-Ape1^Q106A^ | *E. coli* BL21(DE3) strain harboring plasmid pET28a-Ape1^Q106A^ |
| BL21-Ape1^N121A^ | *E. coli* BL21(DE3) strain harboring plasmid pET28a-Ape1^N121A^ |
| BL21-Ape1^S122A^ | *E. coli* BL21(DE3) strain harboring plasmid pET28a-Ape1^S122A^ |
| BL21-Ape1^R123A^ | *E. coli* BL21(DE3) strain harboring plasmid pET28a-Ape1^R123A^ |
| BL21-Ape1^F132A^ | *E. coli* BL21(DE3) strain harboring plasmid pET28a-Ape1^F132A^ |
| **Primer ID** | **Primer Sequence 5′ to 3′ *^b^*** |
| Ape1^41-392^ (F) | GCGGCG***GCTAGC***TCTGCTTTAACAAGTTATGTATCAAAAAAAGATC |
| Ape1^41-392^ (R) | CAATGAGCTCCTAAATTTTTTATAGTAACTAATAATT***CCTAGG***CGCGCG |
| K103A | CTATCCTTTACAGCCT**GCA**TATCAACAAAATC *^c^* |
| Y104G | CCTTTACAGCCTAAA**GGT**CAACAAAATCTTAATC *^c^* |
| Q105A | CTTTACAGCCTAAATAT**GCA**CAAAATCTTAATCTTG *^c^* |
| Q106A | CAGCCTAAATATCAA**GCA**AATCTTAATCTTGTTT *^c^* |
| N121A | CTTTGAAATTTTA**GCC**TCAAGAAATCCTGC *^c^* |
| S122A | CTTTGAAATTTTAAAC**GCA**AGAAATCCTGCTAATGC *^c^* |
| R123A | GAAATTTTAAACTCA**GCA**AATCCTGCTAATG *^c^* |
| F132A | GCTGGACATAAT**GCT**CCACTAGGTGGG *^c^* |

*^a^*Source of strains are as previously described (Ha *et al.*, 2016).

*^b^*Restriction enzyme digestion sites are in bold italics. The three nucleotides of the variant residue are in bold. The nucleotide that is different to the native sequence is underlined.

*^c^*Primers contain 5’ phosphate.

Table S2. HADDOCK calculation statistics of CjApe1-PG complex

| **Docking experiment** | **CjApe1-*O*-acetylated PG** | | | | | | |
| --- | --- | --- | --- | --- | --- | --- | --- |
| **AIR** *^a^* |  | | | | | | |
| Ape1 active residues *^b^* | 73,77,79,82,86,87,90,91,101,103,104,105,106,108,111,112,113,114,121,123,236,237,239,241,272,273,276,277,323,325,328,365,367,369,370,371,372,373,376,380 | | | | | | |
| Ape1 passive residues *^b^* | 62,88,92,110,115,116,117,118,119,122,124,125,126,127,128,147,149,151,153,154,156,244,245,246,274,275,278,279,280,281,285,312,313,318,320,322,326,329,332,333,347,350,359,361,362,366,377,384,392 | | | | | | |
| *O*-acetylated PG active residues *^b^* |  | | | | | | |
| *O*-acetylated PG passive residues *^b^* | 1,2,3,4,5,6 | | | | | | |
| **Cluster** *^c^* |  | | | | | | |
| Cluster no. | Cluster 1 | Cluster 2 | Cluster 3 | Cluster 4 | Cluster 5 | Cluster 6 | Cluster 7 |
| HADDOCK score | -69.2 ± 4.3 | -57.2 ± 3.2 | -50.9 ± 9.3 | -60.0 ± 7.0 | -45.8 ± 10 | -50.7± 8.9 | -54 ± 10.4 |
| Average RMSD (Å) between structures in the cluster | 0.5 ± 0.2 | 0.6 ± 0.2 | 0.4 ± 0.1 | 0.5 ± 0.2 | 0.4 ± 0.2 | 0.4 ± 0.2 | 0.3 ± 0.2 |
| Number of structures in the cluster | 113 | 36 | 11 | 11 | 6 | 6 | 5 |
| Buried surface area (Å^2^) | 1346 ± 54 | 1316 ± 106 | 1281 ± 63 | 1393 ± 57 | 1292 ± 50 | 1327 ± 74 | 1296 ± 21 |
| E_intermolecular_ (kcal/mol) | 131 ± 68 | 148 ± 52 | 230 ± 72 | 162 ± 39 | 128 ± 44 | 200 ± 50 | 130 ± 63 |
| E_non-bonded_ (kcal/mol) | -120 ± 22 | -135 ± 21 | -79 ± 16 | -124 ± 20 | -109 ± 24 | -119 ± 11 | -100 ± 11 |
| E_van der Waals_ (kcal/mol) | -45 ± 4 | -42 ± 4 | -44 ± 5 | -42 ± 3 | -39 ± 8 | -41 ± 4 | -34 ± 4 |
| E_electrostatic_ (kcal/mol) | -75 ± 24 | -93 ± 20 | -35 ± 16 | -82 ± 21 | -71 ± 18 | -79 ± 11 | -66± 8 |
| E_ambiguous intermolecular restraints_ (kcal/mol) | 251 ± 52 | 282 ± 47 | 309 ± 68 | 286 ± 29 | 237 ± 37 | 319 ± 54 | 230 ± 72 |
| Number of AIR violations | 6.1 ± 1.3 | 6.8 ± 1.3 | 7.3 ± 1.8 | 6.8 ± 0.9 | 5.8 ± 1.1 | 8.3 ± 1.4 | 5.4 ± 1.6 |

*^a^*The AIR residues were defined based on the binning interface predicted from CPORT server using Ape1structure and sequence as input information.

*^b^*The number for Ape1 corresponds to the amino acid number in the native sequence. The number 1, 3, 5 and 2, 4, 6 for corresponds to the residues GlcNAc and MurNAc, respectively. Residue 3 is *O*-acetylated MurNAc.

*^c^*In each cluster, the energetic statistics were calculated over the 10 structures with the best HADDOCK scores.

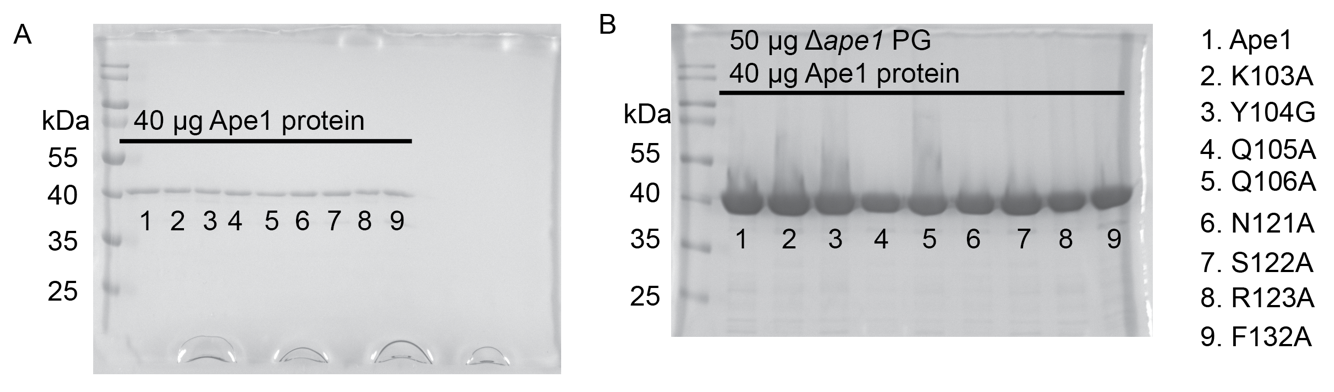

Figure S1. ∆*ape1* PG pull-down assay. 40 µg CjApe1 and variant proteins were incubated in the absence (A) or presence (B) of ∆*ape1* PG in 250 µl buffer (50 mM sodium phosphate buffer, 50 mM NaCl, pH 6.5) at 4 °C for 20 min, followed by three times wash with 1 mL of buffer using centrifugation. The pellet was resuspended in 30 µl of 1× SDS dye (1% (w/v) β- mercaptoethanol, 0.02% (w/v) bromophenol blue, 30% (v/v) glycerol, 10% (w/v) SDS, 250 mM Tris pH6.8). 20 µl of the sample was run on a 12% SDS-PAGE. The number labelled on each lane indicates the corresponding protein (shown as text in right side).

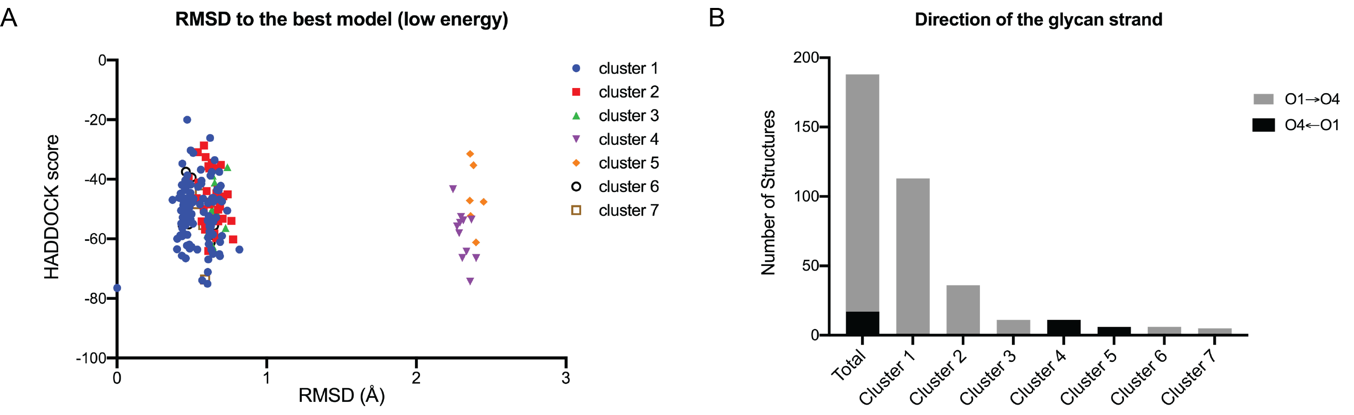

Figure S2. Seven clusters of Ape1-*O*-acetyl hexasaccharide docking models from HADDOCK. (A) The final 188 water-refined models are presented as HADDOCK score in the function of RMSD deviation from the best HADDOCK score model (lowest energy). Models with FCC higher than cut-off (FCC=0.79) are clustered and shown in same color. (B) Clusters 4 and 5 which show large RMSD deviation from the best model contain the glycan strand position in the opposite direction of the best model.

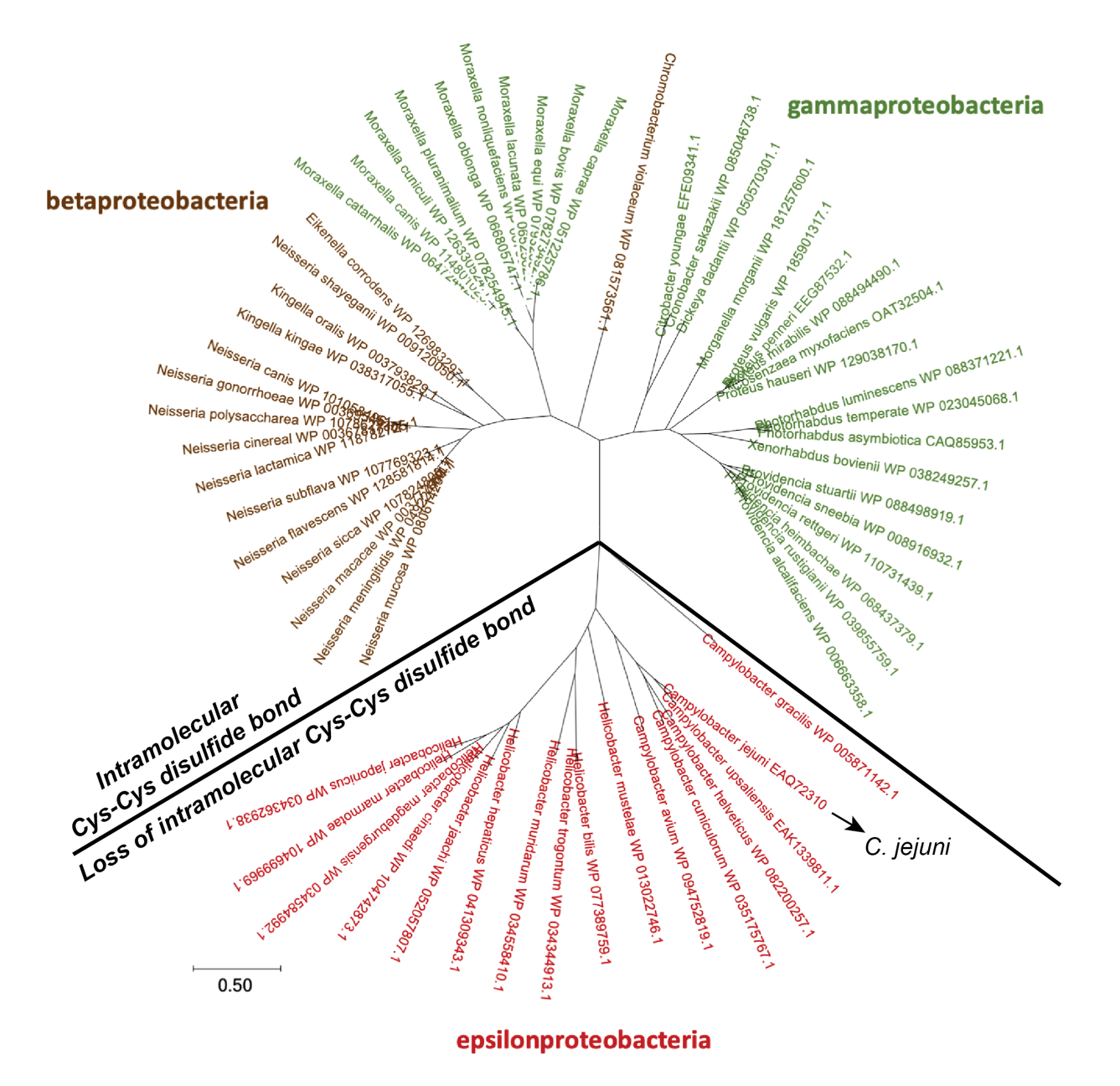
Figure S3. Ape1 phylogenetic tree showing the evolution of the intramolecular disulfide bond of loop SGNHL2. Ape1 homologous sequences (sequence identity: 25%–80%) were searched in the UniRef90 database using ConSurf. Sequences showing ± 20 ammino acids of the length of *C. jejuni* Ape1 (*CJJ81176_0638*) were manually removed. The final 61 sequences were aligned using the MUSCLE alignment algorithm, and a maximum likelihood tree was constructed based on the aligned sequences in MEGA X. Ape1 homologs in betaproteobacteria and gammaproteobacteria contains the intramolecular disulfide bond, whereas Ape1 homologs of epsilonproteobacteria (e.g., *Campylobacter* and *Helicobacter* species) evolved to loss this disulfide bond.

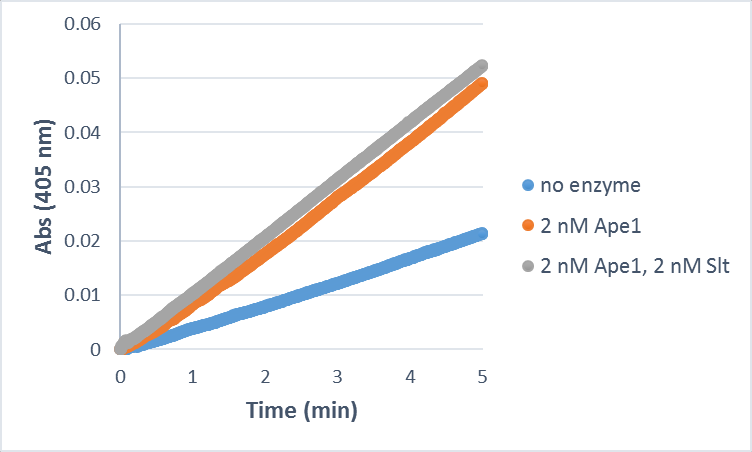

Figure S4. *O*-acetyl esterase activity of CjApe1 and Slt. CjApe1 *O*-acetyl esterase activity with 2 mM *p*NPAc in the presence and absence of equimolar Slt from *C. jejuni* in 50 mM sodium phosphate, 50 mM NaCl, pH 6.5.
